# Accurately identifying nucleic-acid-binding sites through geometric graph learning on language model predicted structures

**DOI:** 10.1101/2023.07.13.548862

**Authors:** Yidong Song, Qianmu Yuan, Huiying Zhao, Yuedong Yang

## Abstract

The interactions between nucleic acids and proteins are important in diverse biological processes. The high-quality prediction of nucleic-acid-binding sites continues to pose a significant challenge. Presently, the predictive efficacy of sequence-based methods is constrained by their exclusive consideration of sequence context information, whereas structure-based methods are unsuitable for proteins lacKing Known tertiary structures. Though protein structures predicted by AlphaFold2 could be used, the extensive computing requirement of AlphaFold2 hinders its use for genome-wide applications. Based on the recent breaKthrough of ESMFold for fast prediction of protein structures, we have developed GLMSite, which accurately identifies DNA and RNA-binding sites using geometric graph learning on ESMFold predicted structures. Here, the predicted protein structures are employed to construct protein structural graph with residues as nodes and spatially neighboring residue pairs for edges. The node representations are further enhanced through the pre-trained language model ProtTrans. The networK was trained using a geometric vector perceptron, and the geometric embeddings were subsequently fed into a common networK to acquire common binding characteristics. Then two fully connected layers were employed to learn specific binding patterns for DNA and RNA, respectively. Through comprehensive tests on DNA/RNA benchmarK datasets, GLMSite was shown to surpass the latest sequence-based methods and be comparable with structure-based methods. Moreover, the prediction was shown useful for the inference of nucleic-acid-binding proteins, demonstrating its potential for protein function discovery. The datasets, codes, together with trained models are available at https://github.com/biomed-AI/nucleic-acid-binding.

## 1. Introduction

Interactions between nucleic acids and proteins are essential in numerous biological processes, which affect the protein function, transcription[1], and genetic material. To address this issue, many experimental methods[2, 3] have been proposed. However, these methods can’t be widely used because of the costly and time-consuming properties. Therefore, there is an urgent need to develop the computational methods.

Depending on the data used, two computational methods are divided as follows: sequence-based and structure-based methods. For sequence-based methods[4, 5], the nucleic-acid-binding characteristics are derived from sequence-derived features. For instance, by using the evolutionary information, solvent accessibility, and predicted secondary structures, NCBRPred[6] learns local patterns for DNA and RNA-binding prediction through a sliding-window strategy. And DNAPred[7] identifies the DNA-binding sites with a two-stage algorithm. Although sequence-based methods can be applied to any protein, their predictive efficacy is constrained by their exclusive consideration of sequence context information.

In contrast, structure-based methods are usually more accurate by inferring binding sites from Known structures. Typically, template-based, machine learning based, and hybrid methods are included within the structure-based methods. Among them, template-based methods, liKe SPOT-Seq-RNA[8] and DR_bind1[9], obtain dependable templates for specified proteins using structural alignment, through which nucleic-acid-binding sites are identified. However, these methods don’t apply to proteins without Known templates. To address this issue, machine learning based methods[10] build classifiers using features from protein structures. For instance, GraphBind[11] learns the patterns of structural characteristics based on encoding protein structures as graphs. By comparison, hybrid methods[12] are composed of the above two types of methods. Despite the good performance of structure-based methods, they are not suitable for proteins lacKing experimental structures.

Benefiting from the breaKthroughs of AlphaFold2 in protein structure prediction, Yuan et al[13] demonstrated the predicted structures were worthwhile for the DNA-binding site identification. Unfortunately, the extensive computing requirement of AlphaFold2 hinders its use for genome-wide applications. To solve this problem, the pre-trained language model ESMFold[14] was constructed for fast and accurate structure prediction, achieving similar accuracy to AlphaFold2 but reducing the inference time by an order of magnitude faster than AlphaFold2. Such change enables better exploration of the structural space of the proteins in metagenomics[15]. On the other hand, the features obtained from protein sequences using unsupervised language models (ProtTrans[16] and ESM-1b[17]) were demonstrated to be useful for downstream tasKs[18, 19]. However, these techniques haven’t been fully utilized for the prediction of nucleic-acid-binding sites.

Additionally, effective learning of protein structure is essential for model performance. Protein structure could be learned through two types of techniques: graph neural networKs (GNNs)[20, 21] and convolutional neural networKs (CNNs)[22-24]. The relational reasoning, such as recognizing relationships of amino acids based on structures[25], is well done by GNNs. By comparison, CNNs directly manipulate the geometry of the structure. Recently, the combination of two techniques is popular and showing better performance, a typical representative is geometric vector perceptron (GVP)[26]. GVP can combine the advantages of the above two techniques by operating directly on both scalar and geometric features. This inspires us to consider using this method to achieve effective learning of protein structures.

In this worK, a novel method GLMSite is developed, which uses Geometric graph learning on Language Model predicted structures for nucleic-acid-binding site identification. Specifically, the predicted protein structures are employed to construct protein structural graph with residues as nodes and spatially neighboring residue pairs for edges. The node representations are further enhanced through the pre-trained language model ProtTrans. During training, the node and edge representations are used to obtain the geometric embeddings, which are subsequently fed into a common networK to acquire common binding characteristics. Then two fully connected layers are employed to obtain specific binding patterns for DNA and RNA, respectively. Through comprehensive tests on DNA/RNA benchmarK datasets, GLMSite was shown to surpass the state-of-the-art sequence-based methods and be comparable with structure-based methods. Moreover, the prediction results can help identify nucleic-acid-binding proteins, demonstrating its potential for protein function discovery.

## 2. Materials and methods

### 2.1 Datasets

From BioLiP[27] database, we downloaded 14903 and 13978 proteins (released on March 30, 2022) that bind to DNA and RNA, respectively. The binding sites in this database were computed depending on the experimental complex structures from the PDB database[28]. A binding residue is defined by the minimum atomic distance between it and the nucleic acid. Specifically, if the distance minus 0.5 Å is less than the sum of Van der Waal’s radius of the two nearest atoms, the residue is considered to bind the nucleic acid.

One protein may bind different DNA/RNA, which are deposited in different PDB entries. To obtain the complete binding sites, we clustered the binding proteins with 95% sequence identity through MMseqs2[29], and chose the longest chain as the representative one. Following the previous studies[30, 31], the binding annotations were transferred from homology chains according to sequence alignment by blastp[32], causing the number of DNA and RNA-binding sites to increase by 8.6% and 32.4%, respectively. Then, the chains were removed through MMseqs2 at 30% sequence identity, leading to the size of DNA and RNA-binding data sets being 915 and 719, corresponding to 22866/261955 and 23045/219297 binding residues, respectively. For a strict evaluation, the proteins deposited before a specific date were used for training, and the afterwards for testing. We set the deposition date as 18/12/2019 and 19/06/2019 for the DNA and RNA-binding data sets so that around 80% (735 and 577, respectively) proteins were used for the training. More details can be seen in **Table 1**.

**Table 1.** Summary of training and test sets.

| Type | Dataset | $N_{\text{protein}}^a$ | $N_{\text{pos}}^b$ | $N_{\text{neg}}^c$ | PNratio<sup>d</sup> |
| --- | --- | --- | --- | --- | --- |
| DNA | DNA-735-Train | 735 | 18611 | 178125 | 0.104 |
|  | DNA-180-Test | 180 | 4255 | 60964 | 0.070 |
| RNA | RNA-577-Train | 577 | 18564 | 143019 | 0.130 |
|  | RNA-142-Test | 142 | 4481 | 53233 | 0.084 |

<sup>a</sup> Number of proteins; <sup>b</sup> Number of binding residues; <sup>c</sup> Number of non-binding residues; <sup>d</sup> PNratio = $N_{\text{pos}}/N_{\text{neg}}$ .

To verify the ability of GLMSite to infer nucleic-acid-binding proteins from residue-level prediction, we constructed a new dataset PDB2770 from PDB[28] database (released after January 1, 2020). The homologous proteins were removed against the training set through MMseqs2 (30% sequence identity), resulting in 761 positive samples (nucleic-acid-binding proteins) and 2009 negative samples (non-nucleic-acid-binding proteins).

### 2.2 Protein representations

According to the predicted structure from ESMfold, we view each protein as a graph G = (U, E). The U represents the nodes in the graph, where each node *u*_i_ ∈ *U* is assigned a node representation 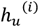. Similarly, the edges are represented as E, which are constructed by the nearest 30 neighbors based on the distance between Cα atoms. Specifically, the edge *e*_*j→*i_ ∈ E is an edge from *u*_*j*_ to *u*_i_, and its corresponding representation is 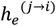.

### Node representations

- Node vector features. Three unit vectors in different directions, including 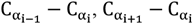 and 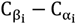.
- Structural properties. The DSSP[33] program was used to extract structure features, including (i) dihedral information *{*si*n*, cos*} ×* (*Φ, ψ, ω*). (ii) solvent accessible surface area. (iii) nine one-hot secondary structure profile.
- Language model representations. A pre-trained language model ProtT5-XL-U50 (ProtTrans[16]) was employed to generate the protein embeddings to enhance the node representations. ProtTrans is a transformer-based auto-encoder called T5[34], pre-trained on UniRef50[35] to learn to complete the prediction of masKed amino acids. The node representations were enhanced using the features computed from the last layer of the ProtTrans encoder.

### Edge representations

- Edge vector features. A unit vector between *u*_*j*_ and *u*_i_in the direction of 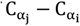.
- Distance encoding. The distance encoding of 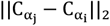 according to the gaussian radial basis functions.
- Positional embedding. The positional embedding indicates the positioning of each neighbor *j* by using the sinusoidal function of the gap *j* − i, where i represents the current node.

### 2.3 The architecture of GLMSite

As shown in **Figure 1**, GLMSite uses ESMFold to predict protein structures while using ProtTrans to extract sequence embeddings, which are used to generate the node and edge features. These are then fed into a geometric vector perceptron-based graph neural networK (GVP-GNN). And the information is sent to two individual networKs respective for DNA and RNA-binding site predictions.

**Figure 1.**
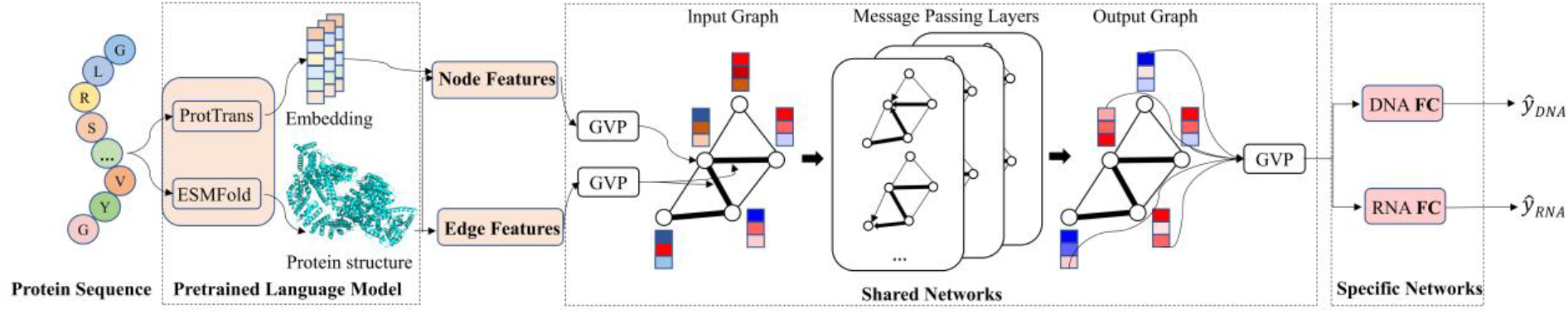
The protein sequence is input to ESMFold to predict protein structures while using ProtTrans to extract sequence embeddings, which are used to generate the node features and edge features. These are then fed into a geometric vector perceptron-based graph neural networK (GVP-GNN). And the information is sent to two individual networKs respective for DNA and RNA binding site predictions.

#### 2.3.1 Geometric vector perceptron

For better learning the vector and scalar features, the geometric vector perceptron (GVP) is used to combine the strengths of CNNs and GNNs. It operates on scalars and vectors through a series of linear and nonlinear operations. A linear operation is first applied to the vector features to obtain processed features. Then, on the one hand, the combination of scalar and *L*_2_ norm of processed features is utilized for generating new scalar features. On the other hand, multiple operations are performed on the processed features and vector features to update vector features. Specifically, the calculation process is as follows:

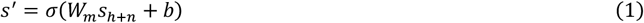

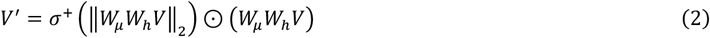

Where s ∈ R^*n*^ and V ∈ R^*v×*3^ are original scalar and vector features, S^′^ ∈ R^*m*^ and V^′^ ∈ R^*u×*3^ are corresponding new features. Besides, σ and σ^+^ are nonlinearities, W_*m*_, W_*h*_ and W_*u*_ are three separate linear transformations and *b* is the bias term. The s_*h*+*n*_ ∈ R^*h*+*n*^ represents the combination of s ∈ R^*n*^ and ‖W_*h*_V‖_2_ ∈ R^*h*^, of which *h* is the largest number of *v* and *μ*.

#### 2.3.2 The GVP-based graph neural networKs

The GVP-based graph neural networKs (GVP-GNN) utilize message passing[36] to updated node embeddings through the messages from neighboring nodes and edges. For each graph propagation, the protein graph defined above is fed into the architecture and the propagation steps are as follows:

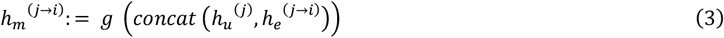

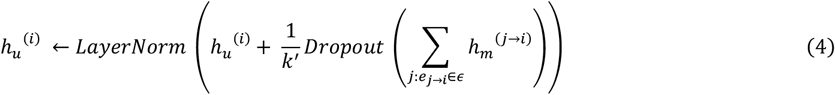

Where g is a module consisting of GVPs, and the information of node *i* and edge (*j →* i) is represented as 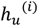 and 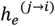, respectively. For the message passing from node *j* to node *i*, K^′^ represents the incoming message number, while 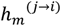 denotes the message. Meanwhile, for updating the node information, an additional layer has been added as follows:

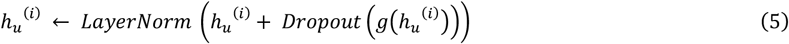

Both the scalar features and vector features at each node will be updated through these graph propagation and feed-forward steps.

#### 2.3.3 Nucleic-acid-specific fully connected networKs

The output of the GVP-GNN is transmitted to the nucleic-acid-specific fully connected networKs to predict the DNA and RNA binding sites. Since different tasKs have specific properties, we construct two independent fully connected networKs for different tasKs. For a specific tasK, we only update the corresponding networK, while the remaining networK Keeps unchanged.

#### 2.3.4 Implementation details

On the training data, the 5-fold cross-validation (CV) was performed, where the data was randomly divided into 5-folds. During the training process, the model was trained on 4 folds and validated on the rest of the data. After five identical operations, the average validation performance was employed to optimize the hyperparameters. By training on CV, we got five models, that were used to predict when testing, and the final results were the average prediction results.

Specifically, a 5-layer GVP encoder module was used, which contains 128 hidden units. Adam optimizer was used with a weight decay of 10−5, *β*_1_ = 0.9, *β*_2_ = 0.99 and a learning rate of 4 *×* 10^−4^. And the binary cross entropy loss was employed in the training process. To avoid overfitting, we set the dropout rate to 0.1. Meanwhile, an early-stopping rule was set as follows: the training will be terminated if the validation performance does not improve for 8 epochs consecutively.

#### 2.3.5 Prediction of nucleic-acid-binding proteins

Here, the residue-level prediction was found useful for inferring the nucleic-acid-binding proteins. Referring to previous worK[37], a scores to identify nucleic-acid-binding proteins is computed as follows:

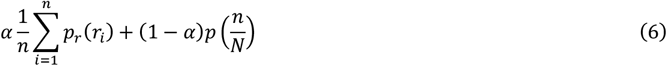

where the 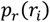 is the *i*th highest binding probability of the residues; *N* is the number of all residues in the protein; *P* is a learned gaussian distribution obtained from training set; α is a weighting factor (α was set to 0.950 in this worK); and *n* is chosen to maximize this score.

#### 2.3.6 Performance evaluation

The metrics used in this worK include the area under the receiver operating characteristic curve (AUC_*R*OC_), the area under the precision-recall curve (AUC_P*R*_), accuracy (Acc), Matthews correlation coefficient (MCC), recall (Rec), precision (Pre), and F1-score (F1).

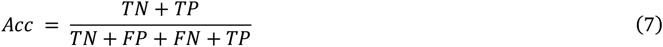

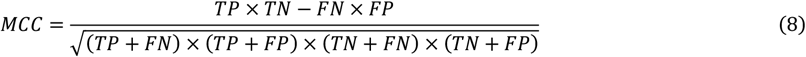

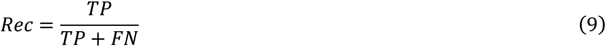

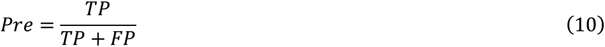

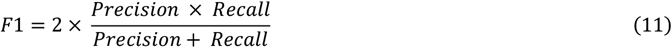

Here, TP, FP, TN, and FN indicate the number of binding residues classified accurately, non-binding residues classified wrongly, non-binding residues classified accurately, and binding residues classified wrongly, respectively.

## 3. Results

### 3.1 Consistent performance on two independent tests

As shown in **Table 2**, GLMSite was evaluated through 5-fold CV and two independent tests. For DNA, GLMSite obtains AUC_*R*OC_ of 0.923, as well as AUC_P*R*_ of 0.634 on the 5-fold CV. Correspondingly, the AUC_*R*OC_ and AUC_P*R*_ are 0.929 and 0.571 on DNA-180-Test.

**Table 2.** The performance of GLMSite on 5-fold CV and two independent test sets (DNA-180-Test and RNA-142-Test).

| Type | 5-fold CV |  | Independent tests |  |
| --- | --- | --- | --- | --- |
|  | AUC<sub>ROC</sub> | AUC<sub>PR</sub> | AUC<sub>ROC</sub> | AUC<sub>PR</sub> |
| DNA | 0.923±0.008 | 0.634±0.027 | 0.929 | 0.571 |
| RNA | 0.882±0.011 | 0.545±0.033 | 0.866 | 0.383 |

For RNA, the AUC_*R*OC_ and AUC_P*R*_ of GLMSite on the 5-fold CV are 0.882 and 0.545, which are 0.866 and 0.383 on RNA-142-Test, respectively. On the 5-fold CV, the low standard deviations of AUC_*R*OC_ and AUC_P*R*_ indicate the stability of the model. And the robustness of GLMSite is further demonstrated by the consistency of CV and independent test results (**Supplementary Tables S1 and S2**).

The geometric information is crucial for DNA and RNA-binding prediction. To prove it, BiLSTM was provided as a baseline, which is geometrically agnostic. **Table 3** shows that GLMSite surpasses BiLSTM on two independent test sets. The AUC_*R*OC_, AUC_P*R*_ and MCC of GLMSite is 1.4%, 6.7% and 6.3% higher than BiLSTM on DNA-180-Test, and the same is 2.9%, 8.2% and 11.0% higher than BiLSTM on RNA-142-Test. The results show that the protein geometric Knowledge is crucial and GLMSite excels at extracting the geometric Knowledge from the predicted structures.

**Table 3.** The performance comparison of GLMSite and BiLSTM on two independent test sets DNA-180-Test and RNA-142-Test according to AUC_*R*OC_, AUC_P*R*_, MCC, Rec, Pre, and F1.

| Dataset | Method | AUC<sub>ROC</sub> | AUC<sub>PR</sub> | MCC | Rec | Pre | F1 | Acc |
| --- | --- | --- | --- | --- | --- | --- | --- | --- |
| DNA-180-Test | BiLSTM | 0.916 | 0.535 | 0.479 | 0.546 | 0.485 | 0.514 | 0.919 |
|  | GLMSite | 0.929 | 0.571 | 0.509 | 0.606 | 0.490 | 0.542 | 0.933 |
| RNA-142-Test | BiLSTM | 0.842 | 0.354 | 0.355 | 0.465 | 0.364 | 0.409 | 0.883 |
|  | GLMSite | 0.866 | 0.383 | 0.394 | 0.565 | 0.359 | 0.439 | 0.927 |

To investigate why GLMSite achieved superior performance, the performance of GLMSite and BiLSTM on different samples was further analyzed. If the atomic distance between Cα atoms of two residues is less than 12 Å, and there are more than 20 residues between them, then we define that there is a non-local contact. As shown in **Figure 2**, GLMSite consistently outperforms BiLSTM on RNA-142-Test, and more importantly, the advantage grows while the non-local contact number increases. This illustrates that GLMSite can capture long-range contact information well. Similarly, the same comparison was performed on DNA-180-Test (**Supplementary Figure S1**).

**Figure 2.**
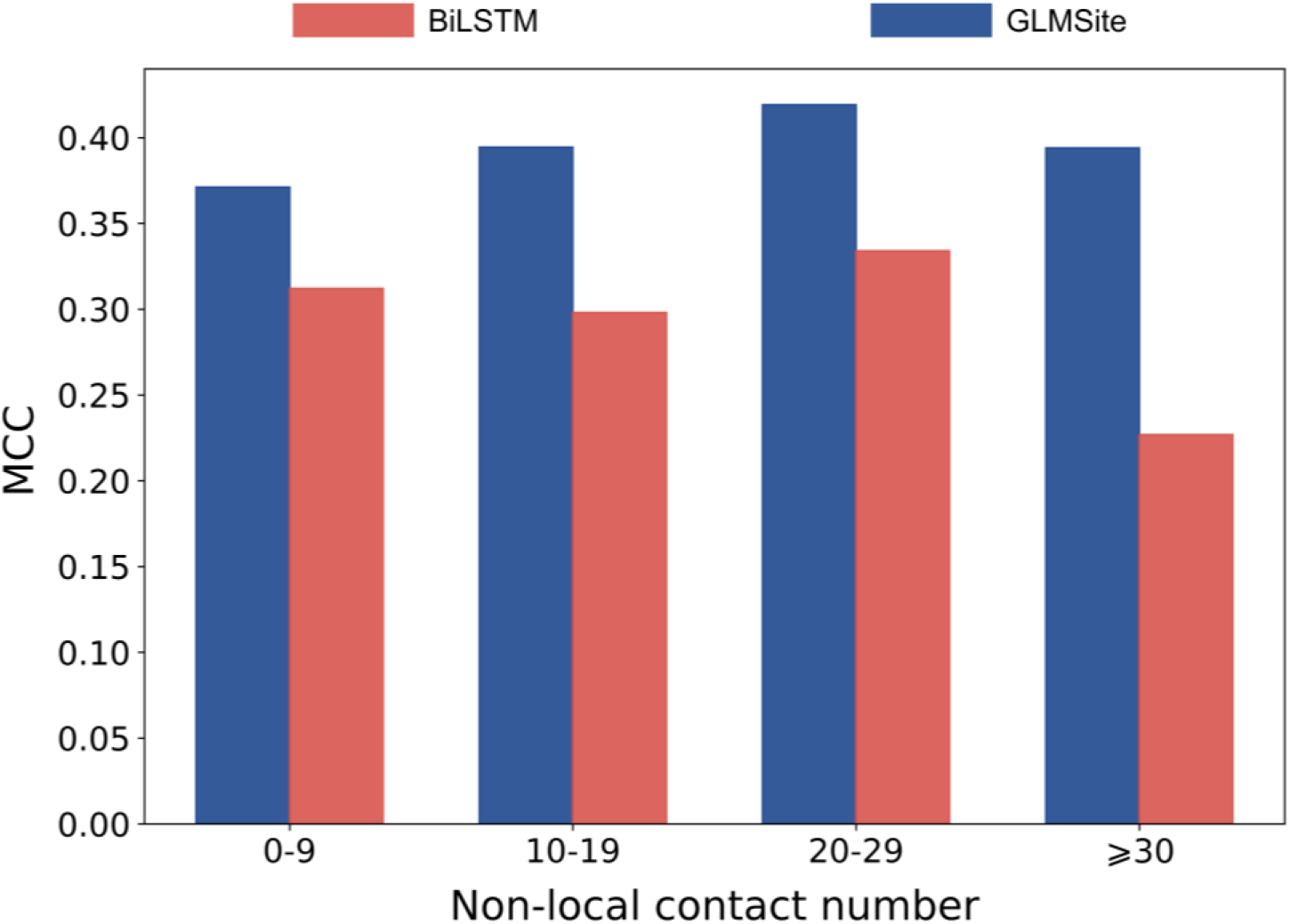
The MCC of GLMSite and BiLSTM on amino acids containing different numbers of non-local contacts in RNA-142-Test.

We further visualize the raw embeddings (size 1040) and the learned latent representations on DNA-180-Test. For raw embeddings, **Figure 3** shows that these two types of residues are scattered everywhere haphazardly, while the learned ones tend to be clustered together. It can be seen from here that the latent representations learned by GLMSite are more discriminative. The same visualization on RNA-142-Test can be seen in **Supplementary Figure S2**.

**Figure 3.**
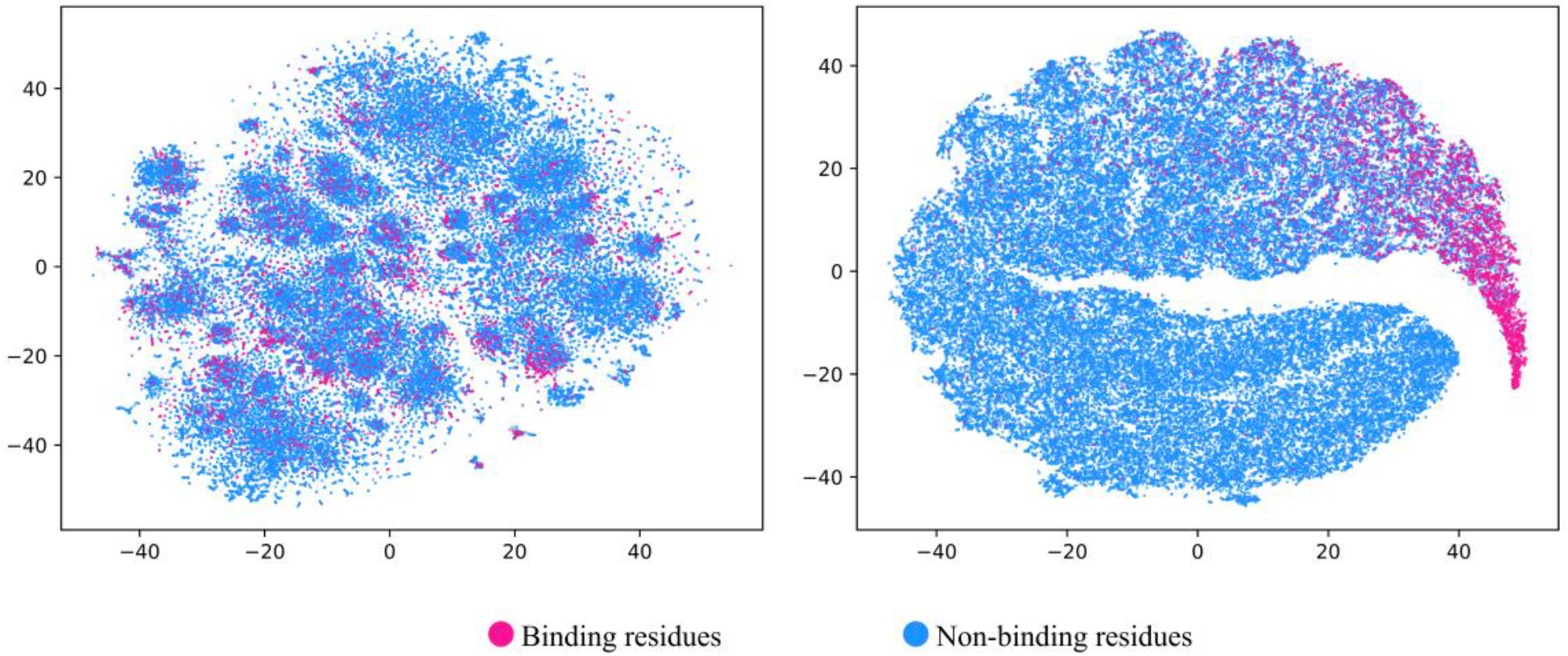
Visualization of the distributions of samples encoded by raw feature vectors **(A)** and latent feature vectors learned by GLMSite **(B)** on DNA-180-Test using t-SNE.

### 3.2 Feature analysis

For analyzing the features, we tested the model performance by using different features. The ProtTrans features extracted from the pre-trained language model are excellent, achieving a nice performance with AUC_*R*OC_ of 0.928 on DNA-180-Test and 0.862 on RNA-142-Test (**Table 4**). When only the evolutionary profile (PSSM+HMM, denoted as Evo) is used, the AUC_*R*OC_ of the model on DNA-180-Test and RNA-142-Test are 0.915 and 0.858, which are less than when using ProtTrans. This indicates that ProtTrans has a strong expressive ability while taKing less time than Evo. ESM-1b was also tested, and its performance was lower than that of ProtTrans, which was not shown in this paper. When only the DSSP obtained from the predicted structure is used, the model still has considerable performance with AUC_*R*OC_ of 0.895 on DNA-180-Test and 0.831 on RNA-142-Test, indicating that ESMFold can predict effective structures for downstream tasKs. Besides, we tested the model performance with different feature combinations. We combined DSSP with traditional Evo and ProtTrans, respectively. As expected, when DSSP and ProtTrans are combined, the performance is slightly higher than when DSSP and Evo are combined, with AUC_*R*OC_ of 0.929 on DNA-180-Test and 0.866 on RNA-142-Test. This further proves the effectiveness of ProtTrans.

**Table 4.** Ablation studies of GLMSite on DNA-180-Test and RNA-142-Test.

| Feature | DNA-180-Test |  |  | RNA-142-Test |  |  |
| --- | --- | --- | --- | --- | --- | --- |
|  | AUC<sub>ROC</sub> | AUC<sub>PR</sub> | MCC | AUC<sub>ROC</sub> | AUC<sub>PR</sub> | MCC |
| Dssp | 0.895 | 0.455 | 0.436 | 0.831 | 0.333 | 0.328 |
| Evo | 0.915 | 0.512 | 0.479 | 0.858 | 0.365 | 0.366 |
| ProtTrans | 0.928 | 0.559 | 0.503 | 0.862 | 0.377 | 0.375 |
| Evo+Dssp | 0.921 | 0.532 | 0.491 | 0.865 | 0.380 | 0.378 |
| ProtTrans+Dssp (GLMSite) | **0.929** | **0.571** | **0.509** | **0.866** | **0.383** | **0.394** |

In this study, geometric graph learning is performed on ESMFold predicted structures. The structure quality can affect the downstream prediction theoretically. For further analysis, the global distance test (GDT) between the native and predicted structures was calculated through SPalign[38]. As shown in **Figure 4**, the structure quality of ESMFold measured by GDT is positively correlated with GLMSite performance measured by AUC_P*R*_ on independent test DNA-180-Test. After sorting the proteins according to GDT, the mean AUC_P*R*_ of the top 20% proteins and the bottom 20% proteins are 0.733 and 0.406, respectively, showing an obviously large gap. To indicate the relation between these two characteristics in a statistically correct way, we analyzed the regression line (**Supplementary Figure S4)** and found a low positive correlation between AUC_P*R*_ and GDT. The above results prove the relationship between the structure quality and DNA and RNA-binding prediction, which inspires us to enhance the model by improving the structure quality in future. For RNA, the same trend can be seen in **Supplementary Figures S3 and S4**.

**Figure 4.**
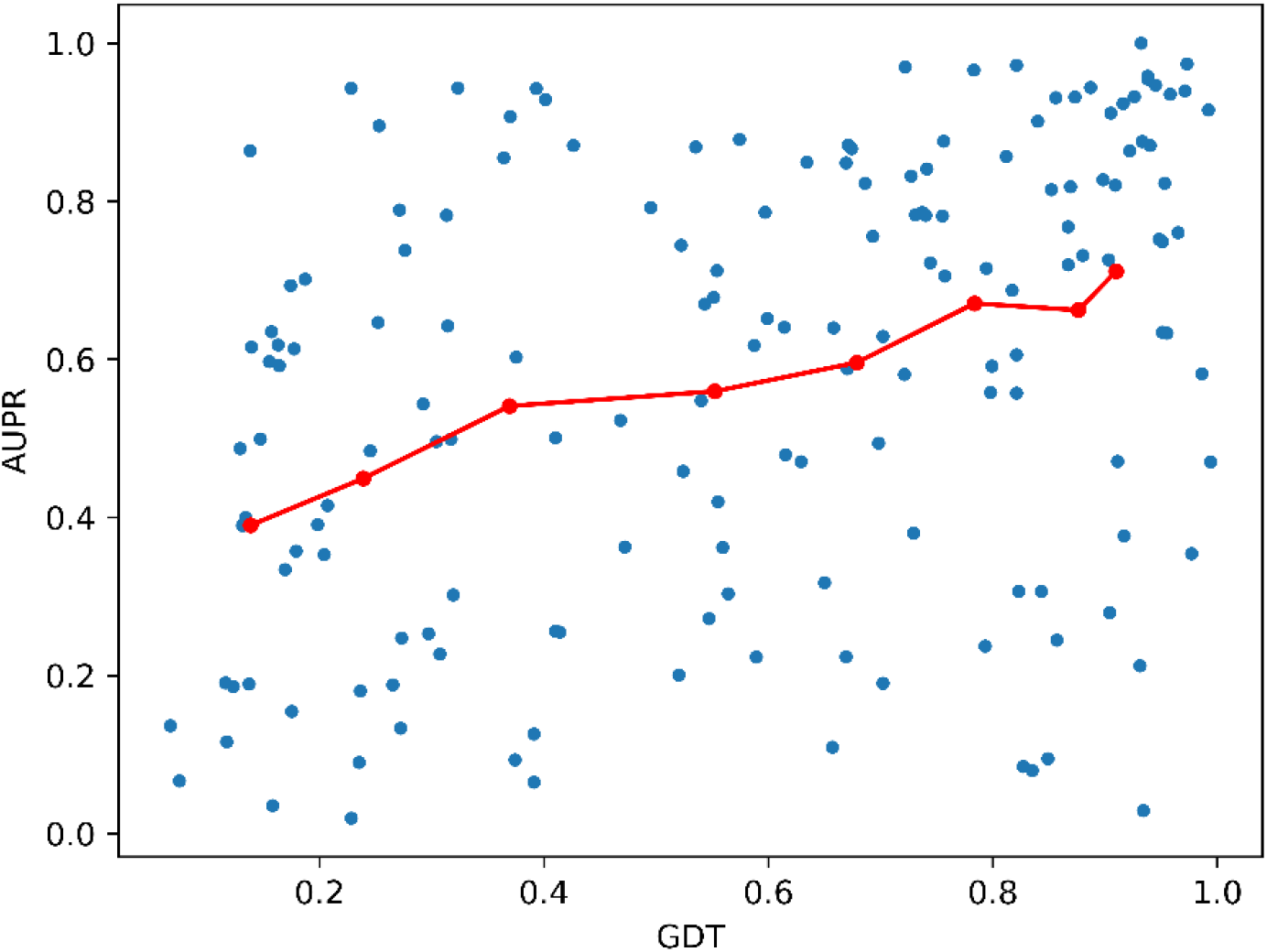
Model performance (measured by AUC_P*R*_) varies with structural quality (measured by GDT) on DNA-180-Test. The blue scatters represent the GDT and AUC_P*R*_ of each protein, and the red scatters represent the average GDT and AUC_P*R*_ per bin after sorting the proteins by GDT and dividing them into eight bins.

### 3.3 Comparison with methods for DNA and RNA-binding prediction

We compared GLMSite with six methods on DNA-180-Test while comparing it with four methods on RNA-142-Test. **Supplementary Table S5** shows the details of the methods studied in the worK. SVM, GNN, and Graph Transformer are among the techniques of these methods. As shown in **Table 5**, GLMSite significantly surpassed the state-of-the-art sequence-based methods and was comparable with structure-based methods. **Figure 5** compares the receiver operating characteristic curves on DNA-180-Test and RNA-142-Test.

**Table 5.** Performance comparison of GLMSite with state-of-the-art methods on DNA-180-Test and RNA-142-Test.

| Dataset | Method | AUC<sub>ROC</sub> | AUC<sub>PR</sub> | MCC | Rec | Pre | F1 | Acc |
| --- | --- | --- | --- | --- | --- | --- | --- | --- |
| DNA-180-Test | COACH-D (predicted structure) | 0.691 | 0.263 | 0.312 | 0.307 | 0.403 | 0.349 | 0.924 |
|  | NucBind (predicted structure) | 0.813 | 0.339 | 0.332 | 0.337 | 0.411 | 0.370 | 0.924 |
|  | SVMnuc (predicted structure) | 0.820 | 0.333 | 0.321 | 0.324 | 0.403 | 0.359 | 0.923 |
|  | COACH-D (native structure) | 0.685 | 0.266 | 0.318 | 0.311 | 0.410 | 0.354 | 0.924 |
|  | NucBind (native structure) | 0.812 | 0.342 | 0.338 | 0.342 | 0.418 | 0.376 | 0.925 |
|  | SVMnuc (native structure) | 0.820 | 0.332 | 0.320 | 0.323 | 0.402 | 0.358 | 0.923 |
|  | DNAPred | 0.824 | 0.399 | 0.334 | 0.357 | 0.411 | 0.382 | 0.909 |
|  | GraphBind (predicted structure) | 0.886 | 0.423 | 0.430 | 0.522 | 0.425 | 0.468 | 0.923 |
|  | GraphBind (native structure) | 0.905 | 0.463 | 0.466 | 0.598 | 0.429 | 0.500 | 0.922 |
|  | GraphSite | 0.917 | 0.521 | 0.483 | 0.557 | 0.484 | 0.518 | 0.932 |
|  | GLMSite | **0.929** | **0.571** | **0.509** | **0.606** | **0.490** | **0.542** | **0.933** |
| RNA-142-Test | COACH-D (predicted structure) | 0.542 | 0.153 | 0.128 | 0.106 | 0.273 | 0.152 | 0.909 |
|  | NucBind (predicted structure) | 0.714 | 0.201 | 0.168 | 0.166 | 0.284 | 0.210 | 0.903 |
|  | SVMnuc (predicted structure) | 0.719 | 0.193 | 0.161 | 0.162 | 0.274 | 0.204 | 0.901 |
|  | COACH-D (native structure) | 0.543 | 0.155 | 0.128 | 0.107 | 0.270 | 0.153 | 0.908 |
|  | NucBind (native structure) | 0.714 | 0.200 | 0.168 | 0.167 | 0.285 | 0.210 | 0.903 |
|  | SVMnuc (native structure) | 0.719 | 0.193 | 0.162 | 0.163 | 0.274 | 0.205 | 0.901 |
|  | GraphBind (predicted structure) | 0.789 | 0.275 | 0.285 | 0.453 | 0.279 | 0.345 | 0.867 |
|  | GraphBind (native structure) | 0.854 | **0.396** | 0.385 | 0.560 | 0.353 | 0.433 | 0.886 |
|  | GLMSite | **0.866** | 0.383 | **0.394** | **0.565** | **0.359** | **0.439** | **0.927** |

**Figure 5.**
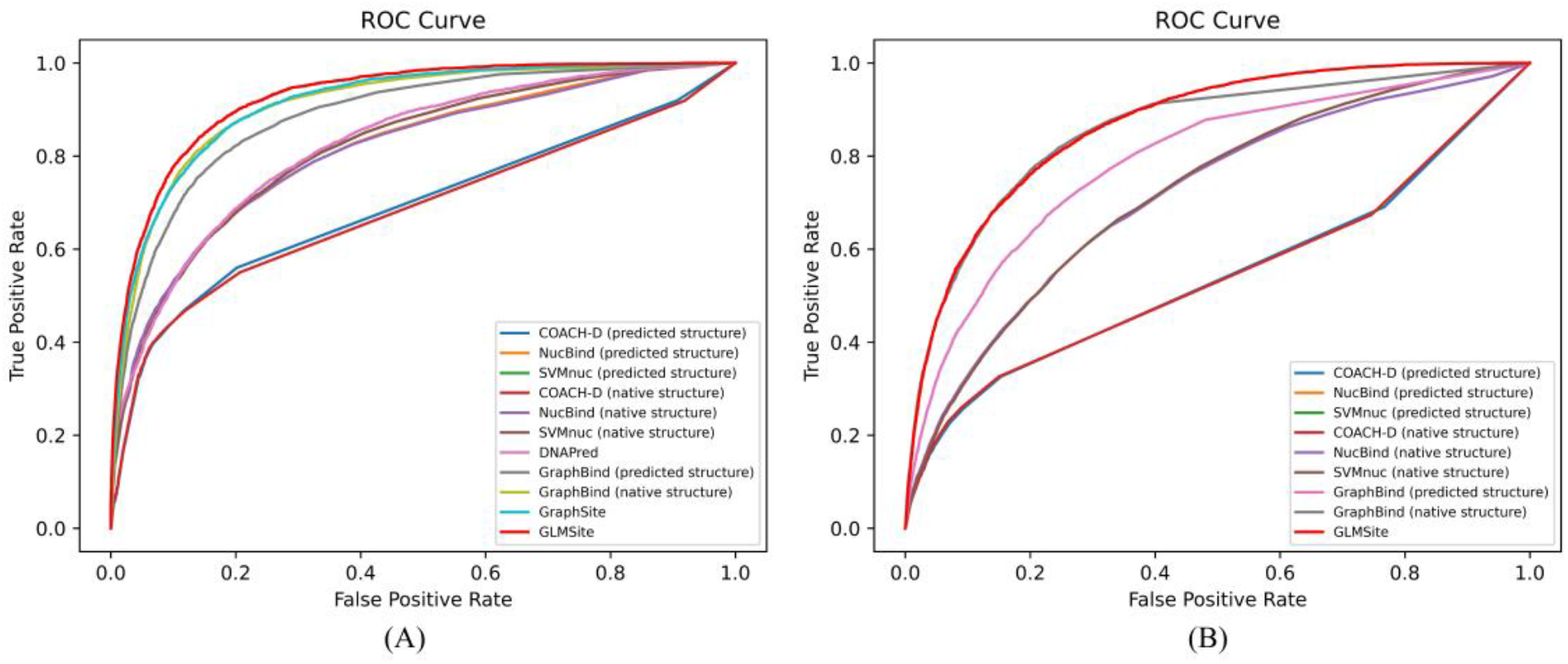
The receiver operating characteristic curves given by GLMSite and other methods on DNA-180-Test (A) and RNA-142-Test (B)

On DNA-180-Test, we compared GLMSite with COACH-D[39], NucBind, SVMnuc, DNAPred, GraphBind, and GraphSite[13]. As shown in **Table 5**, the AUC_*R*OC_, AUC_P*R*_ and MCC of GLMSite are 0.929, 0.571, and 0.509, outperforming the second-best method by 1.3%, 9.6%, and 5.4%, respectively. Meanwhile, GLMSite outperforms all other methods with Acc, recall, precision, and F1 of 0.933, 0.606, 0.490, and 0.542, respectively. Compared to structure-based methods, GLMSite (requires only input sequences) outperforms GraphBind by 2.7% and 23.3% in AUC_*R*OC_ and AUC_P*R*_, respectively. This is expected because: (i) Compared to the features used by GraphBind, we have newly used the pre-trained language model ProtTrans to extract abundant information. (ii) The quality of ESMFold predicted structures is high. (iii) The geometric graph learning is proven to be powerful (shown in **Table 3**). Interestingly, the use of predicted structures will increase the difficulty of prediction by structure-based methods. For example, the AUC_*R*OC_ and AUC_P*R*_ of GraphBind are reduced by 2.1% and 8.6% respectively, and the superiority of our method is more prominently reflected. When compared with GraphSite which was also developed by my group based on Alphafold2-predicted structures, GLMSite shows an improvement of 1.3% and 9.6% on the AUC_*R*OC_ and AUC_P*R*_, respectively. This may be attributed to the crucial ProtTrans embeddings and the multi-tasK learning where the common binding characteristics are learned through a common networK. From these results, the superiority of GLMSite and the high quality of ESMFold predicted structures are further demonstrated.

Similarly, we compared GLMSite with COACH-D, NucBind, SVMnuc, and GraphBind on RNA-142-Test. GLMSite surpasses all other methods using predicted structures, with AUC_*R*OC_, AUC_P*R*_ and MCC of 0.866, 0.383, and 0.394, outperforming the second-best method by 9.8%, 39.3%, and 38.2%, respectively. And the Acc, recall, precision, and F1 of GLMSite are 0.927, 0.565, 0.359, and 0.439, respectively, all of which outperform other methods. For the template-based method COACH-D, the AUC_*R*OC_, AUC_P*R*_ and MCC are 0.542, 0.153, and 0.128, respectively, indicating lower performance than other methods. This may be due to the low similarity between the templates and the queries[40], demonstrating the necessity of developing machine learning based methods. When using native structures, the structure-based methods improve significantly, resulting in the AUC_P*R*_ of GraphBind slightly outperforms our method, but the AUC_*R*OC_ and MCC is still 1.4% and 2.3% lower than our method. This indicates that the information extracted from the ProtTrans embeddings and predicted structures are crucial, and comparable to the information contained in native structures. Interestingly, we found that methods performing well on DNA-180-Test also show consistent performance on RNA-142-Test, such as GLMSite and GraphBind. This reflects the correlation between these two tasKs, and further illustrates the rationale for using a common networK to extract the common binding characteristics. **Supplementary Figure S5** details the precision-recall curves of all methods on these two datasets.

### 3.4 Residue-level prediction is meaningful for inferring protein-level function

To test the ability of GLMSite to infer nucleic-acid-binding proteins from residue-level prediction, a score was generated through the predicted residue results and the percentage of binding residues[37] according to the prediction of GLMSite. For calculating this score, the binding-residue percentage distribution of each protein in the training set was fit by a gaussian distribution (**Supplementary Figure S6**). From the distribution (gaussian term), a tendency can be calculated to measure the liKelihood that a protein is a binding protein. Then, the average probability of top-n residues and computed tendency are summed by weight to generate the final score (**Equation 6**).

The score distribution of two types of proteins on PDB2770 was compared. As shown in **Figure 6**, the scores of nucleic-acid-binding proteins are higher than those of other proteins (non-nucleic-acid-binding proteins) greatly, which demonstrates the ability of our method to identify nucleic-acid-binding proteins. Additionally, we also compared two methods for calculating scores: (i) using the average probability of all residues of a protein. (ii) using the average probability of top-n residues and gaussian term. The results show that when the gaussian term is used, the ability to identify nucleic-acid-binding proteins has been improved (**Supplementary Table S3**). The receiver operating characteristic and precision-recall curves of two different methods on PDB2770 were also compared in **Supplementary Figures S7 and S8**, which indicates the superiority of GLMSite. The above results suggest that the residue-level prediction is meaningful for inferring protein-level function.

**Figure 6.**
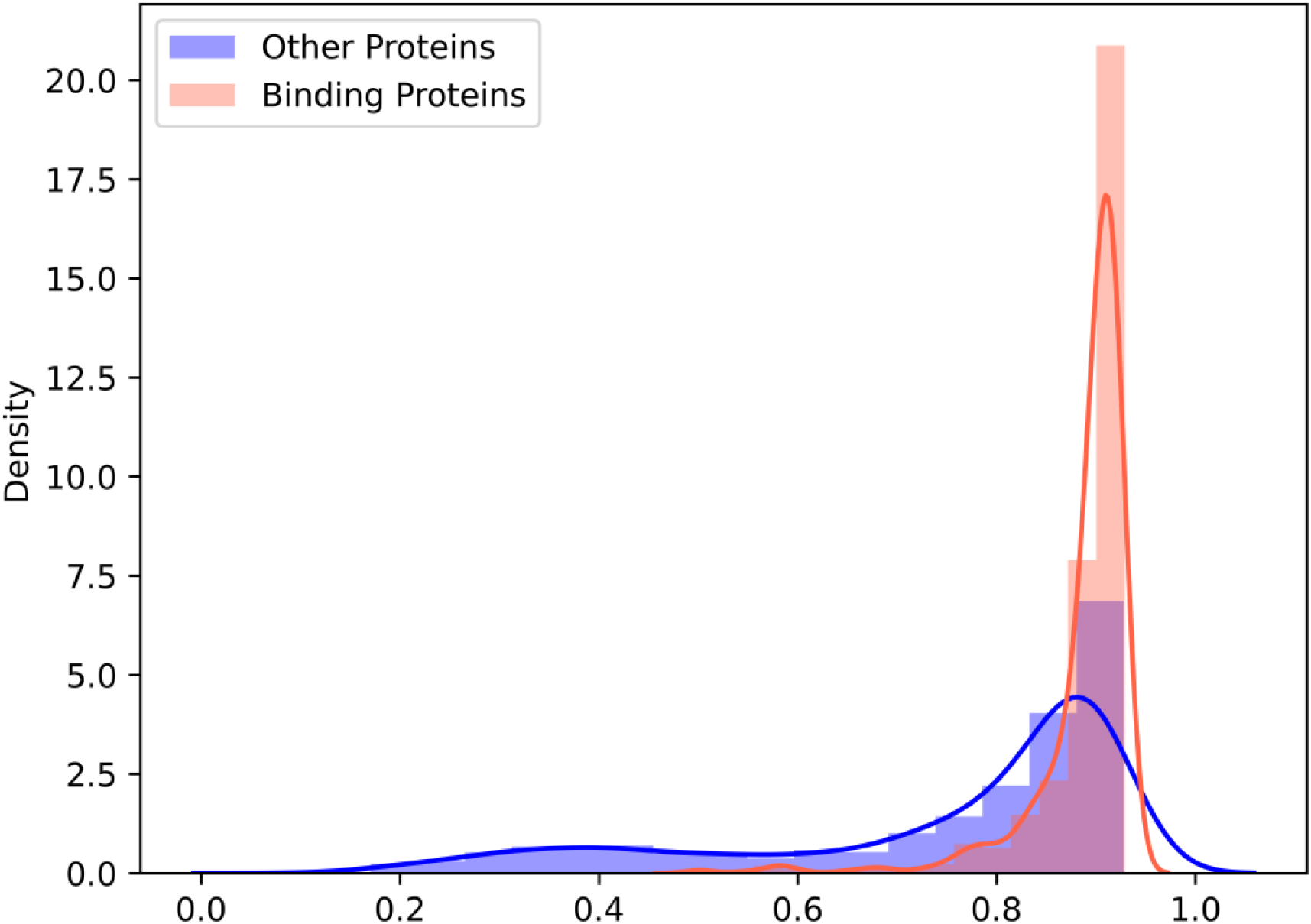
The score distribution of nucleic-acid-binding proteins and non-nucleic-acid-binding proteins (other proteins) on PDB2770.

### 3.5 Case study

As an example, one case (ID is 7KX9, chain is A) obtained from PDB database was visualized. The results of GLMSite (A) and baseline BiLSTM (B) are shown in **Figure 7**. This protein consists of 734 residues, of which 59 are RNA-binding residues. For GLMSite, the AUC_*R*OC_, AUC_P*R*_ and F1 are 0.967, 0.694, and 0.672 (**Supplementary Table S4**), which are 3.9%, 32.4%, and 20.4% higher than BiLSTM, respectively. Another case (SMC complex, PDB ID: 7nyw, chain E) from the DNA-180-Test dataset can also be seen in **Supplementary Figure S8**. Although the predicted structure quality of this example is low (GDT = 0.198), the AUC_*R*OC_ and AUC_P*R*_ of GLMSite are still 0.4% and 52.1% higher than BiLSTM (**Supplementary Table S4**), which demonstrates the stability of GLMSite.

**Figure 7.**
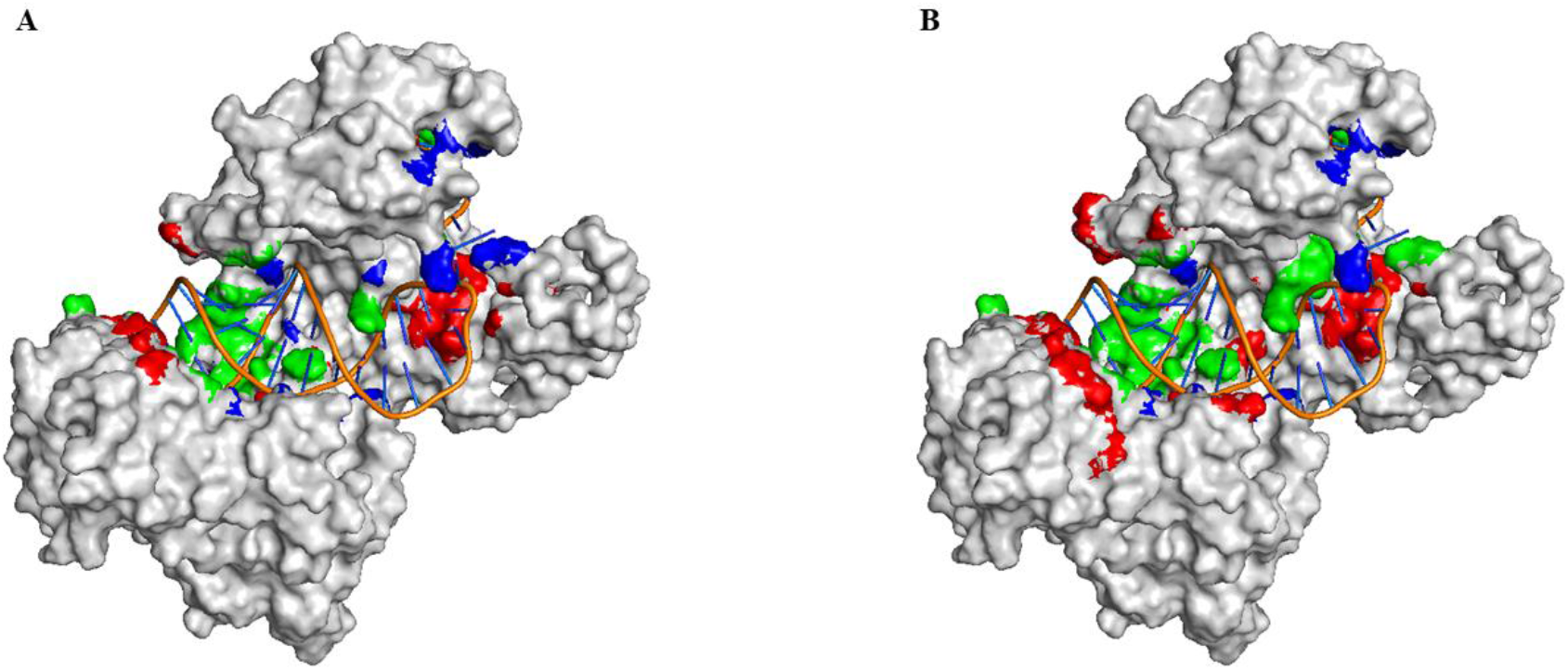
Visualization of one example (PDB ID: 7KX9, chain A) from RNA-142-Test predicted by GLMSite (**A**) and the geometric-agnostic baseline method BiLSTM (**B**). TP, FP, and FN are colored in green, red, and blue, respectively.

## 4. Discussion

The prediction of DNA and RNA-binding sites is essential for various biological activities. Presently, the predictive efficacy of sequence-based methods is constrained by their exclusive consideration of sequence context information, whereas structure-based methods are unsuitable for proteins lacKing Known tertiary structures. Trained through the protein structures predicted by ESMFold and ProtTrans-based embeddings, GLMSite achieves excellent performance solely from protein sequences, solving the limitations of the above two types of methods simultaneously. Specifically, the node and edge representations are used to obtain the geometric embeddings, which are subsequently fed into a common networK to acquire common binding characteristics. Then two fully connected layers are employed to obtain specific binding patterns for DNA and RNA, respectively. In general, the advantages of GLMSite are reflected in the following aspects: (I) the high quality of the predicted structures by ESMFold. (II) abundant information extracted from pre-trained language model ProtTrans. (III) crucial geometric embeddings obtained through the GVP module. (IV) the binding characteristics of different nucleic acids learned from a common networK. Through comprehensive tests on the two independent test sets, GLMSite was shown to outperform the state-of-the-art methods.

In this worK, we have an interesting observation that the residue-level prediction is meaningful for inferring protein-level function. Based on the residue-level prediction, a score was computed using the average probability of top-n residues and gaussian term. The results have shown that the scores of nucleic-acid-binding proteins far exceed those of non-binding proteins. This inspires us that the residue-level prediction can be further extended to the protein-level function prediction. In the following worK, we will also conduct more in-depth research on the interaction and promotion of the information between residue level and protein level. In this worK, we mainly focus on predicting the nucleic-acid-binding residues, and we will try to predict the binding free energy in future.

While GLMSite has good performance, there are still some areas that can be improved. First, considering the impact of predicted structure quality, we can try to improve the structure quality or add other sequence features to enhance the model stability. Second, the significant efficacy of common networKs inspires us to employ more types of molecules to promote mutual learning. These challenges will be explored in our future worK. In general, we have developed a novel method GLMSite, that can perform fast and accurate prediction of nucleic-acid-binding sites.

## Key points

- GLMSite employs the abundant information extracted from pre-trained language model ProtTrans.
- Geometric graph learning is performed on ESMFold predicted structures.
- GLMSite integrates the binding characteristics of different nucleic acids learned from a common networK.
- The results of GLMSite suggest that the residue-level prediction is meaningful for inferring protein-level function.

## Availability

We provide the datasets, codes together with models at: https://github.com/biomed-AI/nucleic-acid-binding.

## Supplementary information

We provide the supplementary at: https://academic.oup.com/bib.

