## Supplementary for "Accurately identifying nucleic-acid-binding sites through geometric graph learning on language model predicted structures" (from another version of this preprint: version 1, 2023-07-15)

Author biography:

**Yidong Song** is a Ph.D. student in the School of Computer Science and Engineering at Sun Yat-sen University. His research interests include protein function prediction, graph neural network, and protein disorder prediction.

**Qianmu Yuan** is a Ph.D. student in the School of Computer Science and Engineering at Sun Yat-sen University. His research interests include deep learning, protein function prediction, and graph neural network.

**Huiying Zhao** is an associate research fellow in the Sun Yat-sen Memorial Hospital at Sun Yat-sen University. Her research interests include protein function, pathogenic gene analysis, and RNA function prediction.

**Yuedong Yang** is a professor in the School of Computer Science and Engineering at Sun Yat-sen University. Currently he focuses on integrating HPC and AI algorithms for biomedical research.

**Supplementary Note 1:** The evolutionary information was used for ablation study. The evolutionary-derived features containing hidden Markov models (HMM) and position-specific scoring matrix (PSSM) were tested for feature ablation studies. HHM profile was obtained from HHblits[1] by aligning the query sequence with the UniClust30[2]. Meanwhile, PSI-BLAST[3] was employed to compute PSSM by searching against UniRef90[4] with an E-value of 0.001 and three iterations.

**Table S1.** The performance of GLMSite on the 5-fold cross-validation (CV) and DNA-180-Test with different hyperparameters. The hyperparameters that are not shown in each row are the same as the final GLMSite model.

| Hyperparameter | Value | 5-Fold CV $\mathrm{AUC}_{\mathrm{ROC}}$ | 5-Fold CV $\mathrm{AUC}_{\mathrm{PR}}$ | DNA-180-Test  $\mathrm{AUC}_{\mathrm{ROC}}$ | DNA-180-Test  $\mathrm{AUC}_{\mathrm{ROC}}$ |
| --- | --- | --- | --- | --- | --- |
| Layers | 3 | 0.923 | 0.611 | 0.922 | 0.530 |
|  | **5** | **0.923** | **0.634** | **0.929** | **0.571** |
|  | 7 | 0.920 | 0.622 | 0.922 | 0.533 |
| Learning rate | $\boldsymbol{4 \times}\mathbf{10}^{\mathbf{-4}}$ | **0.923** | **0.634** | **0.929** | **0.571** |
|  | $4 \times{10}^{-5}$ | 0.917 | 0.614 | 0.915 | 0.516 |
|  | $4 \times{10}^{-6}$ | 0.917 | 0.611 | 0.917 | 0.522 |
| Patience | 6 | 0.922 | 0.624 | 0.923 | 0.536 |
|  | **8** | **0.923** | **0.634** | **0.929** | **0.571** |
|  | 10 | 0.916 | 0.618 | 0.924 | 0.548 |
| Hidden Units | 32 | 0.921 | 0.623 | 0.923 | 0.539 |
|  | 64 | 0.920 | 0.618 | 0.923 | 0.532 |
|  | **128** | **0.923** | **0.634** | **0.929** | **0.571** |
| Dropout | **0.1** | **0.923** | **0.634** | **0.929** | **0.571** |
|  | 0.2 | 0.919 | 0.615 | 0.921 | 0.528 |
|  | 0.3 | 0.918 | 0.610 | 0.919 | 0.519 |

**Table S2.** The performance of GLMSite on the 5-fold cross-validation (CV) and RNA-142-Test with different hyperparameters. The hyperparameters that are not shown in each row are the same as the final GLMSite model.

| Hyperparameter | Value | 5-Fold CV $\mathrm{AUC}_{\mathrm{ROC}}$ | 5-Fold CV $\mathrm{AUC}_{\mathrm{PR}}$ | RNA-142-Test  $\mathrm{AUC}_{\mathrm{ROC}}$ | RNA-142-Test  $\mathrm{AUC}_{\mathrm{PR}}$ |
| --- | --- | --- | --- | --- | --- |
| Layers | 3 | 0.882 | 0.542 | 0.856 | 0.358 |
|  | **5** | **0.882** | **0.545** | **0.866** | **0.383** |
|  | 7 | 0.876 | 0.528 | 0.854 | 0.365 |
| Learning rate | $\boldsymbol{4 \times}\mathbf{10}^{\mathbf{-4}}$ | **0.882** | **0.545** | **0.866** | **0.383** |
|  | $4 \times{10}^{-5}$ | 0.874 | 0.529 | 0.846 | 0.344 |
|  | $4 \times{10}^{-6}$ | 0.875 | 0.533 | 0.845 | 0.335 |
| Patience | 6 | 0.878 | 0.531 | 0.854 | 0.357 |
|  | **8** | **0.882** | **0.545** | **0.866** | **0.383** |
|  | 10 | 0.875 | 0.539 | 0.857 | 0.363 |
| Hidden Units | 32 | 0.880 | 0.541 | 0.859 | 0.367 |
|  | 64 | 0.877 | 0.535 | 0.856 | 0.359 |
|  | **128** | **0.882** | **0.545** | **0.866** | **0.383** |
| Dropout | **0.1** | **0.882** | **0.545** | **0.866** | **0.383** |
|  | 0.2 | 0.877 | 0.534 | 0.852 | 0.349 |
|  | 0.3 | 0.877 | 0.534 | 0.850 | 0.346 |

**Table S3.** The comparison of two different methods for calculating scores on PDB2770, where Method1 represents calculating scores with average probability of all residues of a protein, and Method2 represents calculating scores using the average probability of top-n residues and gaussian term.

| Method | $\mathrm{AUC}_{\mathrm{ROC}}$ | $\mathrm{AUC}_{\mathrm{PR}}$ | MCC | Rec | Pre | F1 |
| --- | --- | --- | --- | --- | --- | --- |
| Method1 | 0.698 | 0.443 | 0.267 | 0.585 | 0.428 | 0.494 |
| Method2 | **0.811** | **0.610** | **0.431** | **0.603** | **0.578** | **0.590** |

**Table S4.** The performance comparison of GLMSite and BiLSTM about the case of 7kx9_A in RNA-142-Test.

| Method | $\mathrm{AUC}_{\mathrm{ROC}}$ | $\mathrm{AUC}_{\mathrm{PR}}$ | MCC | Rec | Pre | F1 | # binding residues | # predicted binding residues | True positive |
| --- | --- | --- | --- | --- | --- | --- | --- | --- | --- |
| BiLSTM | 0.931 | 0.524 | 0.518 | 0.610 | 0.514 | 0.558 | 59 | 70 | 36 |
| GLMSite | **0.967** | **0.694** | **0.643** | **0.678** | **0.667** | **0.672** | 59 | 60 | 40 |

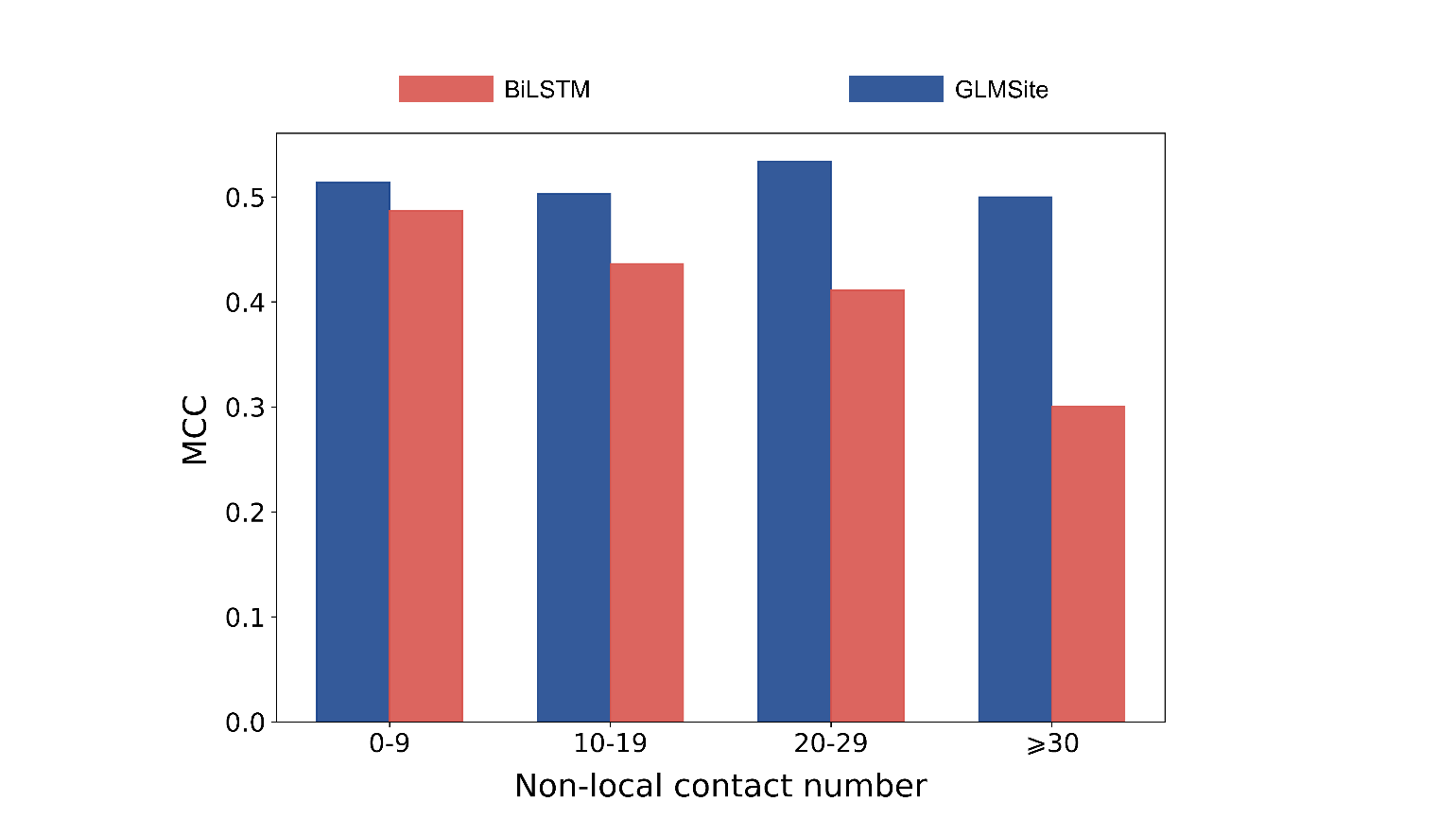

**Figure S1.** The MCC of GLMSite and BiLSTM on amino acids containing different numbers of non-local contacts in DNA-180-Test

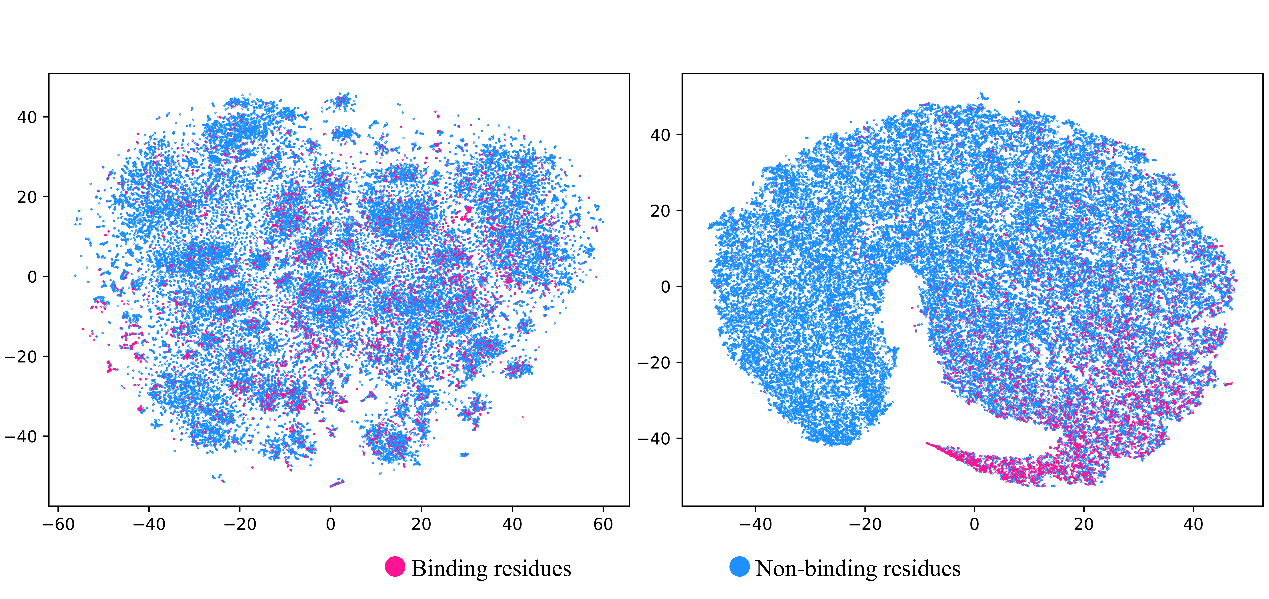

**Figure S2**. Visualization of the distributions of samples encoded by raw feature vectors (A) and latent feature vectors learned by GLMSite (B) on RNA-142-Test using t-SNE.

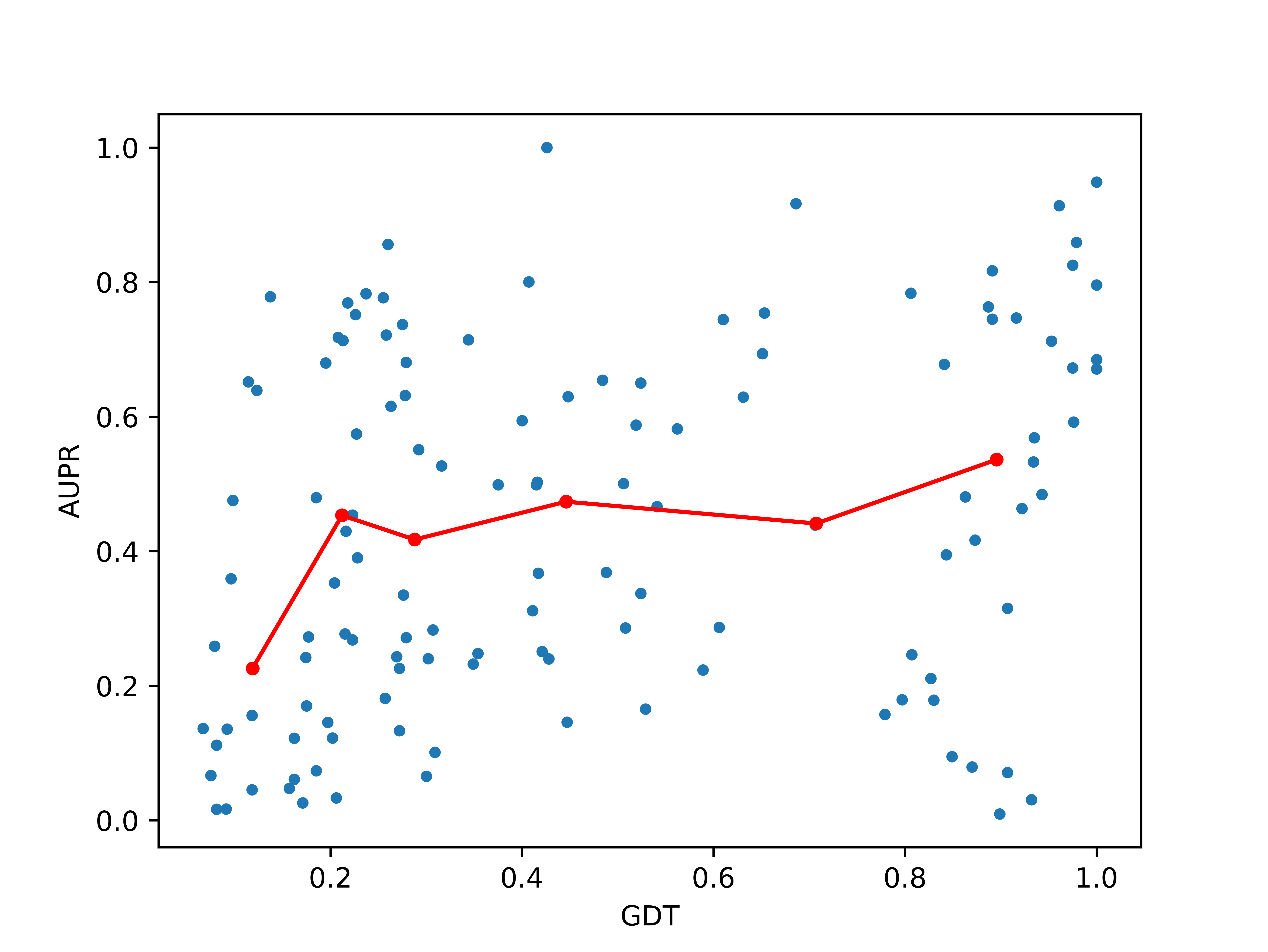

**Figure S3**. Model performance (measured by $\mathrm{AUC}_{\mathrm{PR}}$) varies with structural quality (measured by GDT) on RNA-142-Test. The blue scatters represent the GDT and $\mathrm{AUC}_{\mathrm{PR}}$ of each protein, and the red scatters represent the average GDT and $\mathrm{AUC}_{\mathrm{PR}}$ per bin after sorting the proteins by GDT and dividing them into six bins.

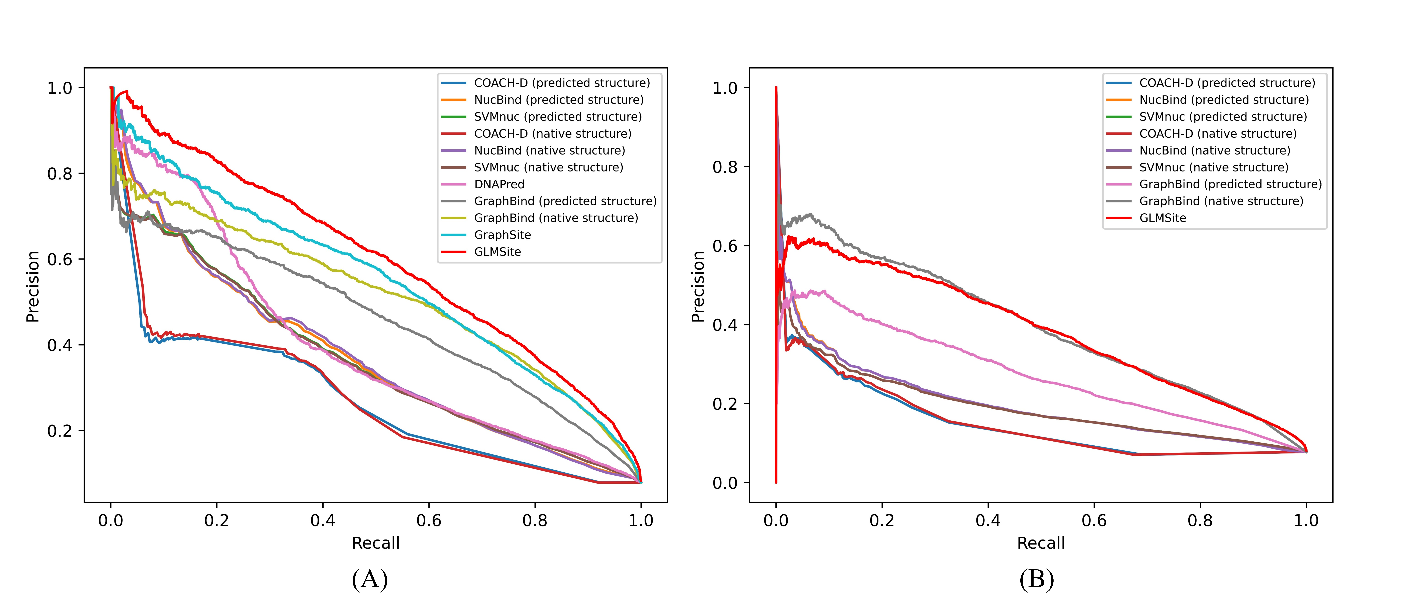

**Figure S4**. The precision-recall curves given by GLMSite and other methods on DNA-180-Test (A) and RNA-142-Test (B)

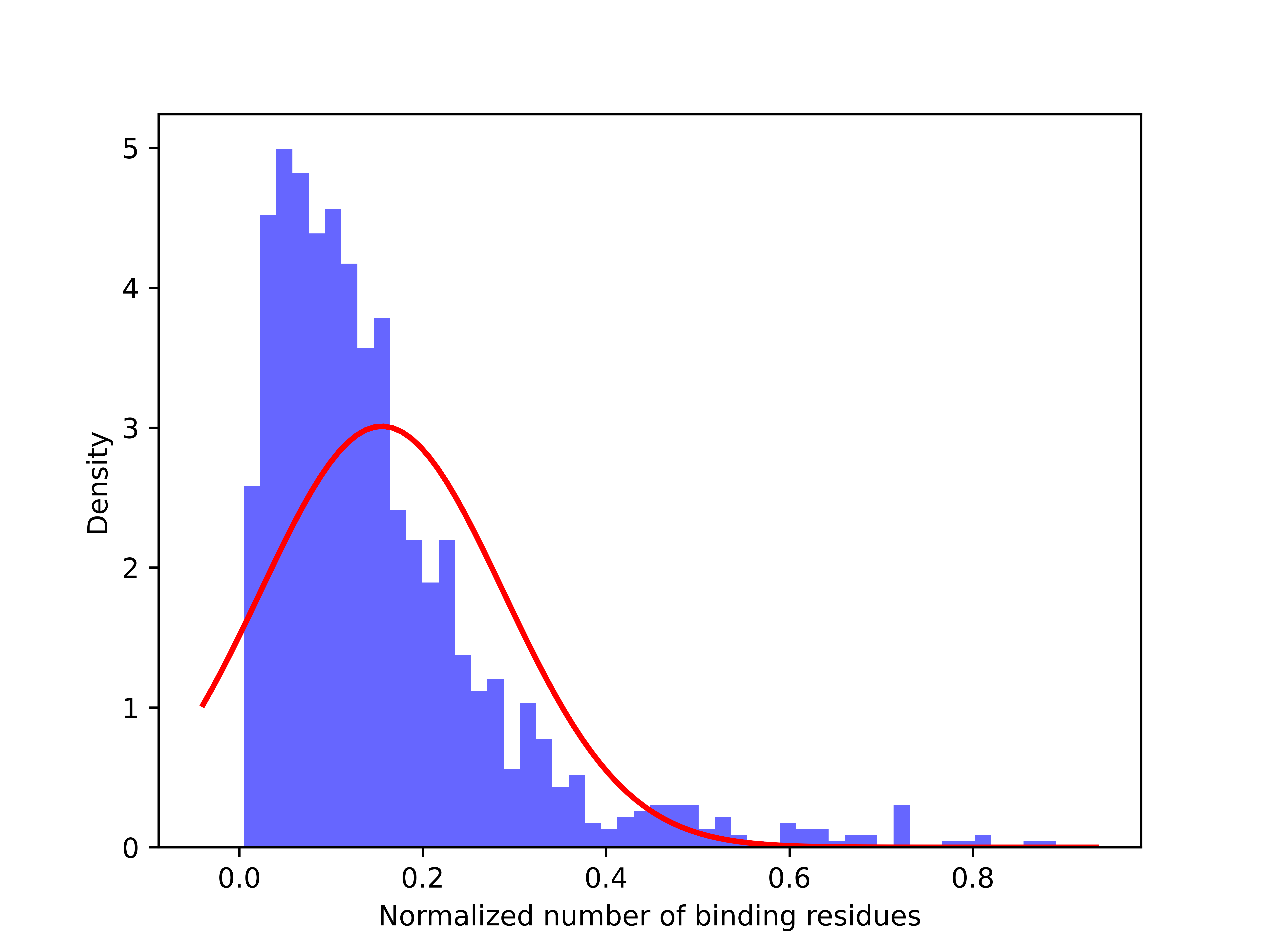

**Figure S5**. The percentage distribution of binding residues in the training set.

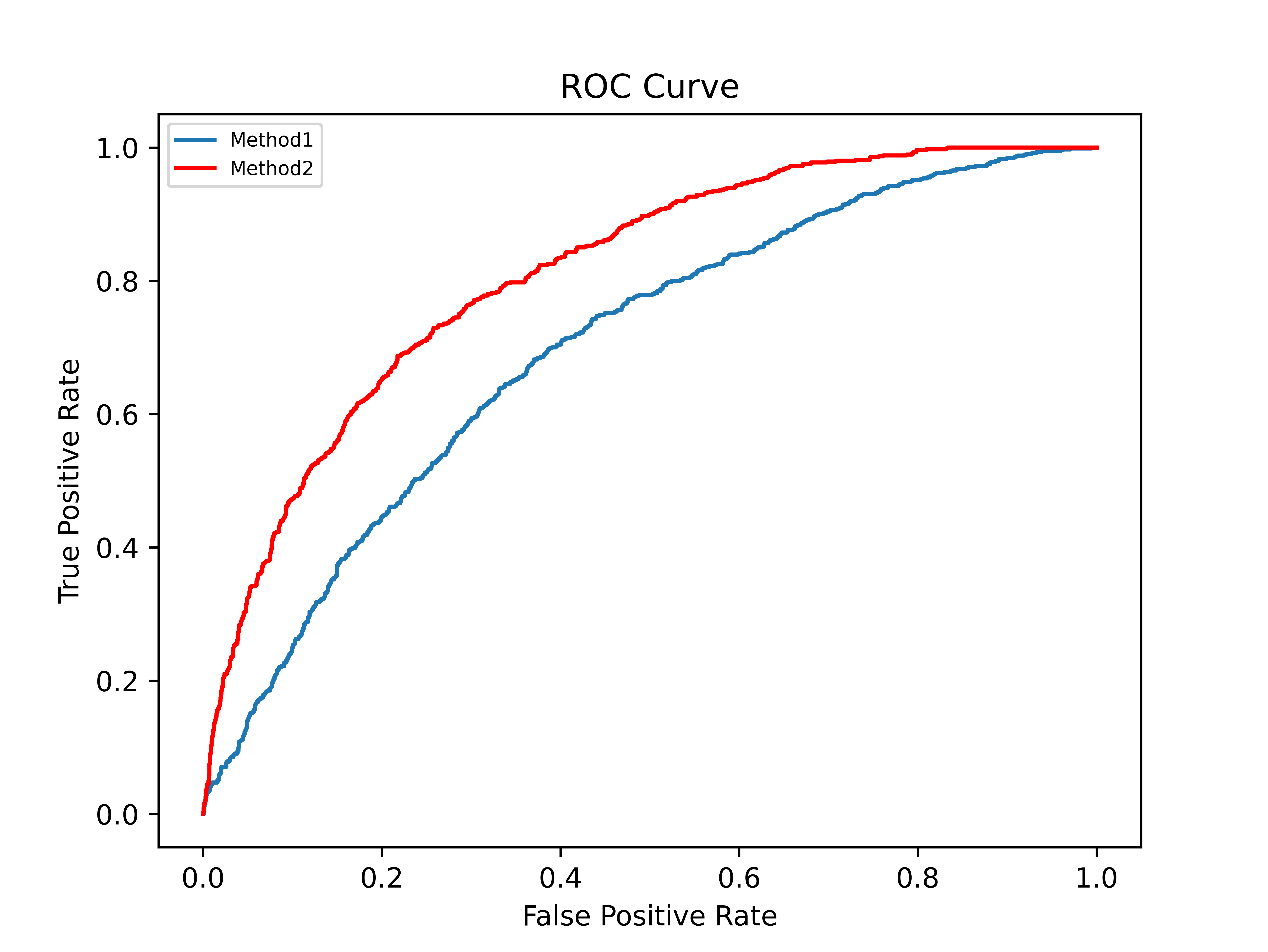

**Figure S6**. The receiver operating characteristic curves given by two methods on PDB2770, where Method1 represents calculating scores with average probability of all residues of a protein, and Method2 represents calculating scores using the average probability of top-n residues and gaussian term.

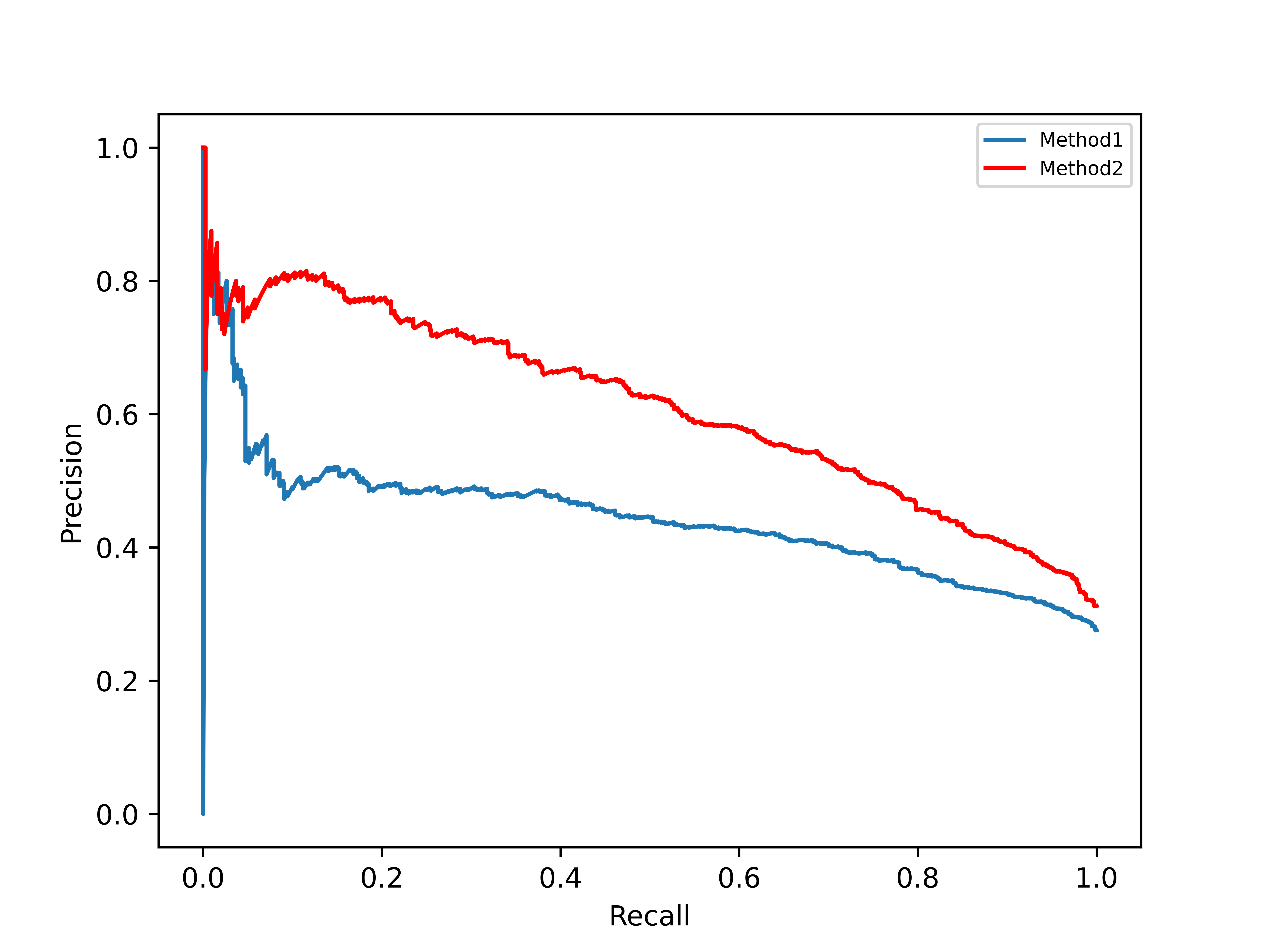

**Figure S7**. The precision-recall curves given by two methods on PDB2770, where Method1 represents calculating scores with average probability of all residues of a protein, and Method2 represents calculating scores using the average probability of top-n residues and gaussian term.
